## Supplemental Figures for "Single-cell and single-nucleus RNA-sequencing from paired normal-adenocarcinoma lung samples provide both common and discordant biological insights"

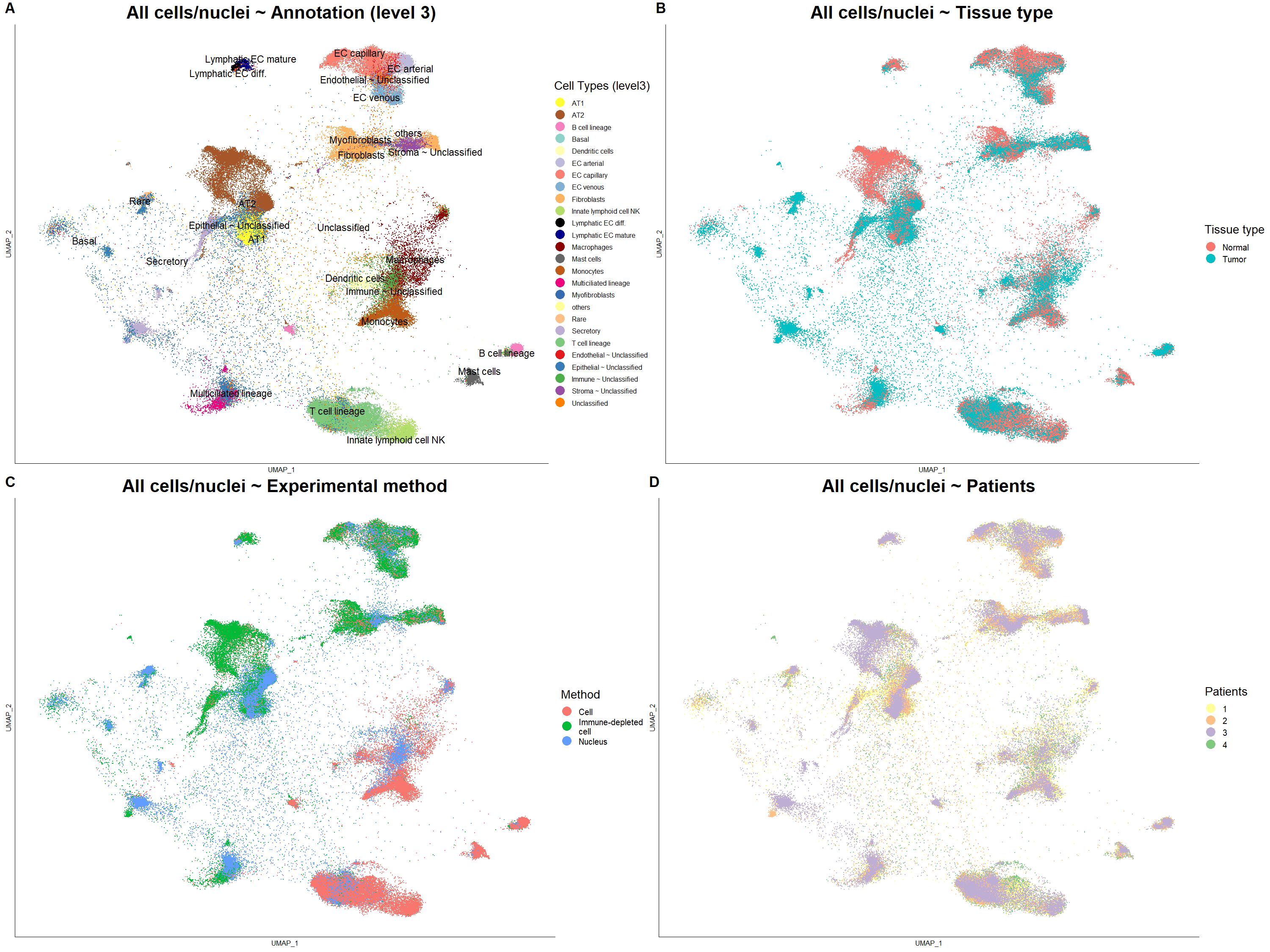

**S1 Fig |** UMAP visualization of all 160,621 cells / nuclei that passed quality control per level 3 annotation (**A**), tissue type (**B**), experimental method (**C**) and patient (**D**).

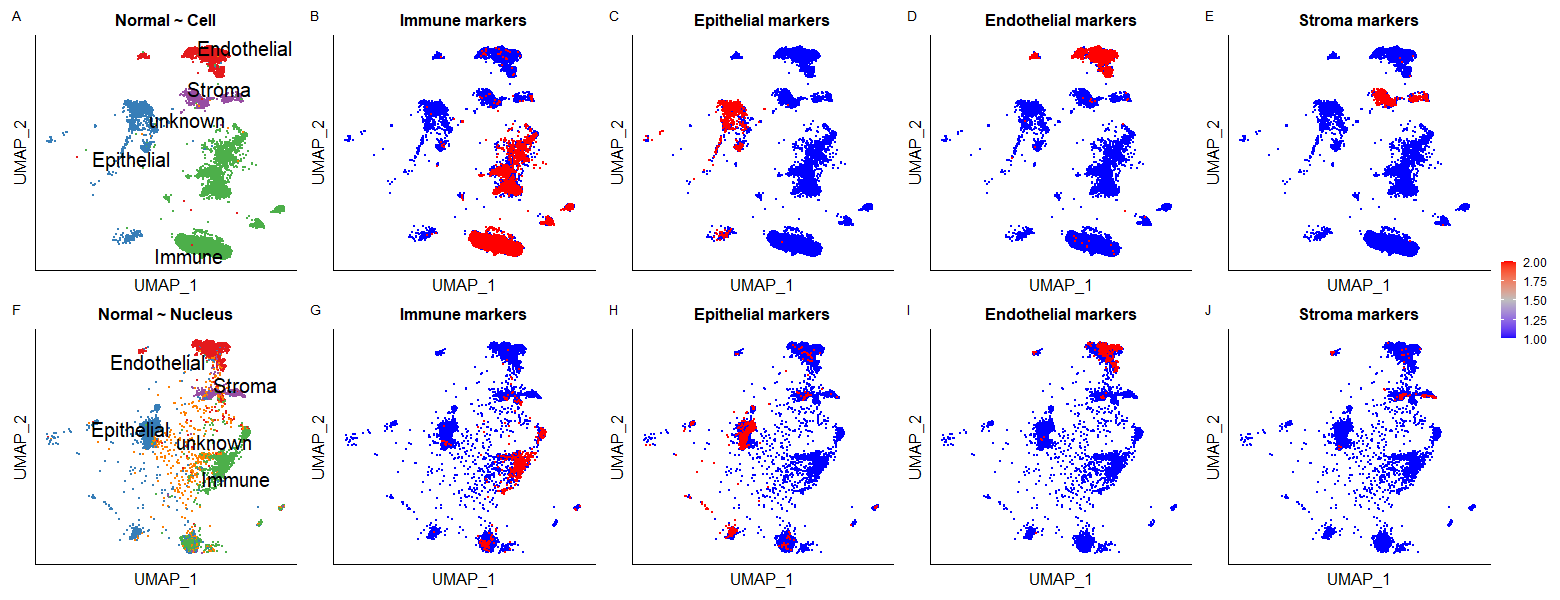

**S2 Fig | UMAPs for the Cell (A) and Nucleus (F) dataset** with coarse level annotations and feature plots according to average expression level of the gene markers defined for each cell type by HLCA (see below), in *Cell* (**B-E)** and *Nucleus* (**G-J**).

Immune-specific gene markers = 'LCP1','CD53','PTPRC','COTL1','CXCR4','GMFG','FCER1G','LAPTM5','SRGN','CD52'

Epithelial-specific gene markers = 'KRT7','PIGR','ELF3','CYB5A','KRT8','KRT19','TACSTD2','MUC1','S100A14','CXCL17'

Endothelial-specific gene markers = 'PTRF','CLDN5','AQP1','PECAM1','NPDC1','VWF','GNG11','RAMP2','CLEC14A'

Stroma-specific gene markers = 'TPM2','DCN','MGP','SPARC','CALD1','LUM','TAGLN','IGFBP7','COL1A2','C1S'

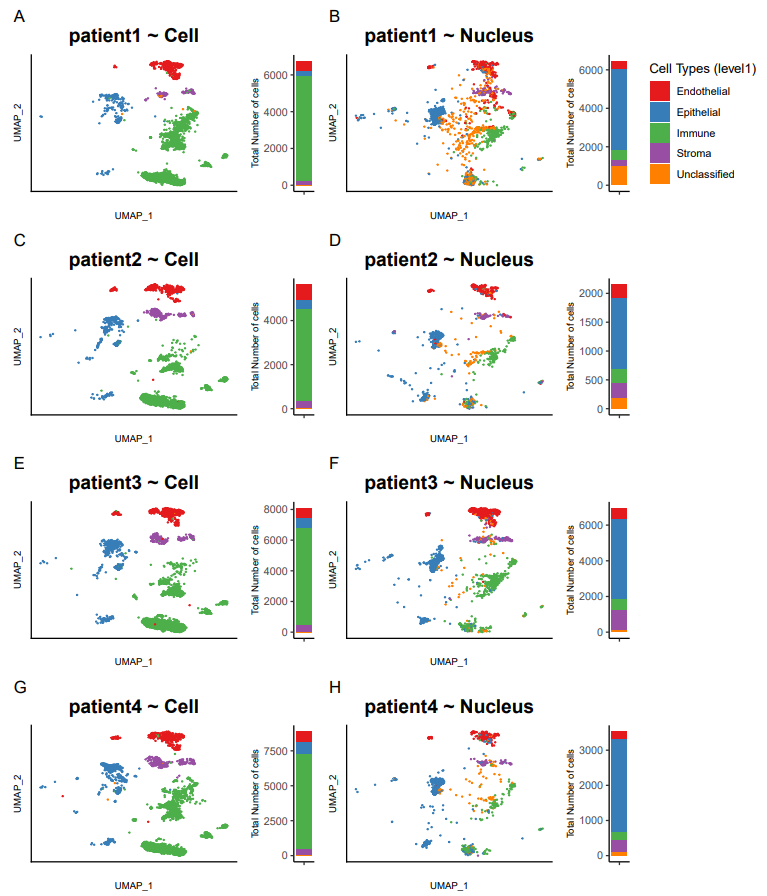

**S3 Fig|** UMAP per patients for Normal samples

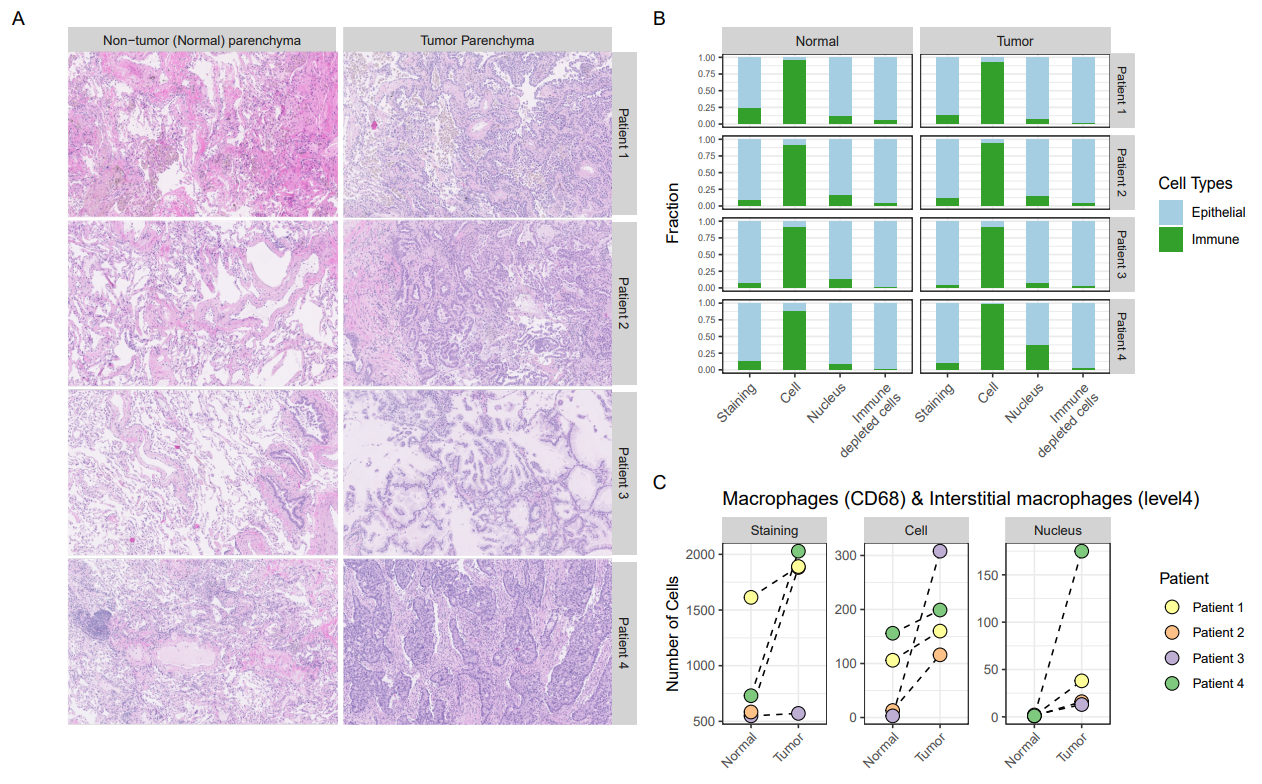

**S4 Fig | A.** Hematoxylin and Eosin staining of Normal and Tumor lung parenchyma used for cell isolation. 100X magnification. **B.** Fraction of Epithelial (AE1/AE3) and Immune (CD45) cells identified through immunohistochemical staining compared to Epithelial and Immune cells (level 1), obtained for the three experimental methods, i.e. *Cell*, *Nucleus* and *Immune depleted cell*. **C.** Number of macrophages (CD68) identified through immunohistochemical staining compared to the most relevant cell type (Interstitial macrophage, level 4) for the *Cell* and *Nucleus* datasets. The *Immune depleted cell* dataset was excluded because the number of macrophages was insufficient.

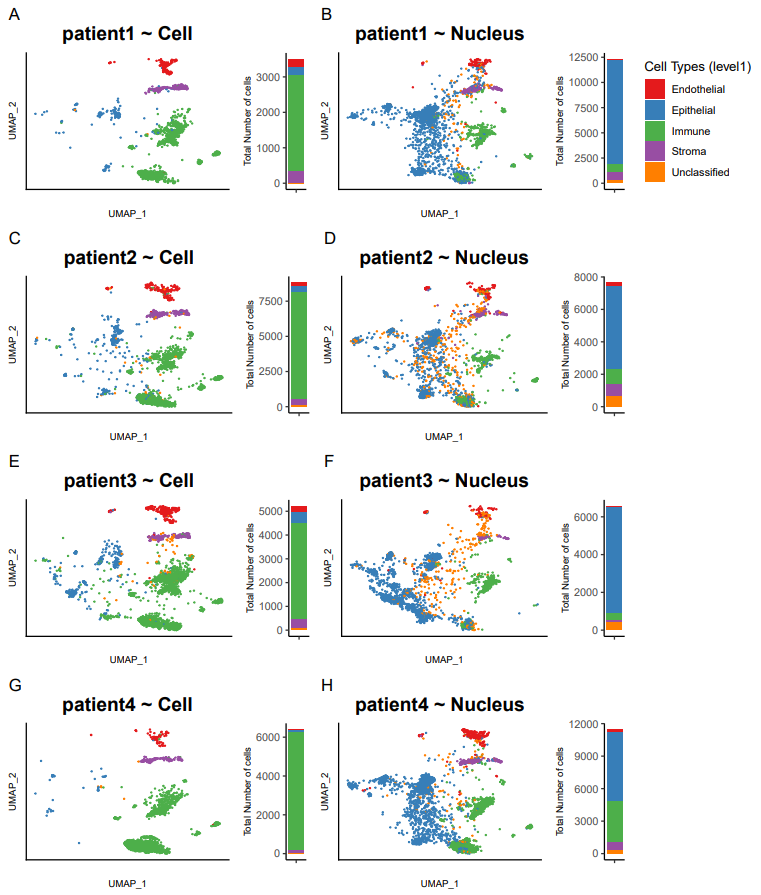

**S5 Fig |** UMAP per patients for Tumor samples

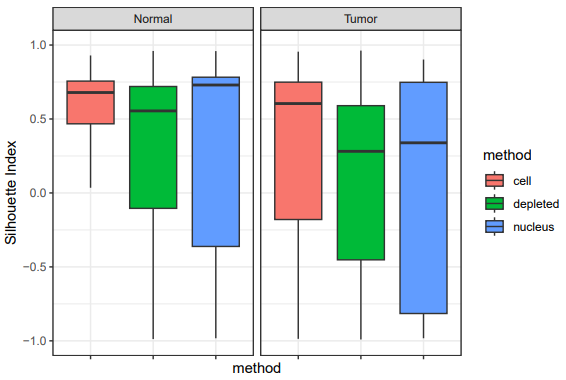

**S6 Fig | Silhouette index to evaluate the goodness of fit of the clustering**. For each cell / nucleus, Silhouette Indices are calculated from the UMAP embeddings and the clusters correspond to a specific cell type (level 3) annotations. Silhouette Index was significantly lower (less structured clusters) for *Tumor* rather than *Normal* samples.

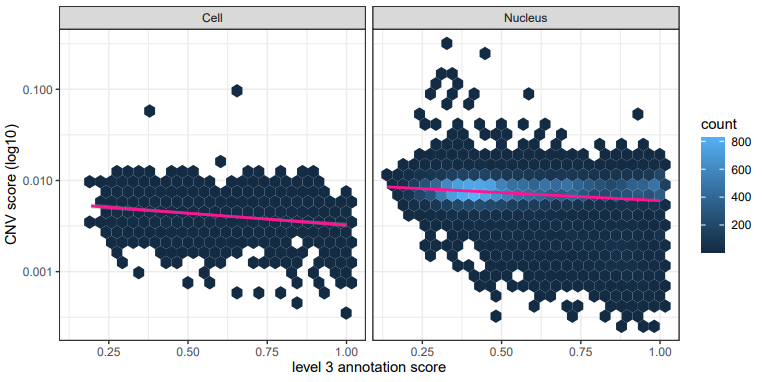

**S7 Fig | Annotation score (level 3) is negatively correlated with CNV score.** Data points were binned (50 hexagonal bins in x-axis * 50 hexagonal bins in y-axis) to reduce overplotting.

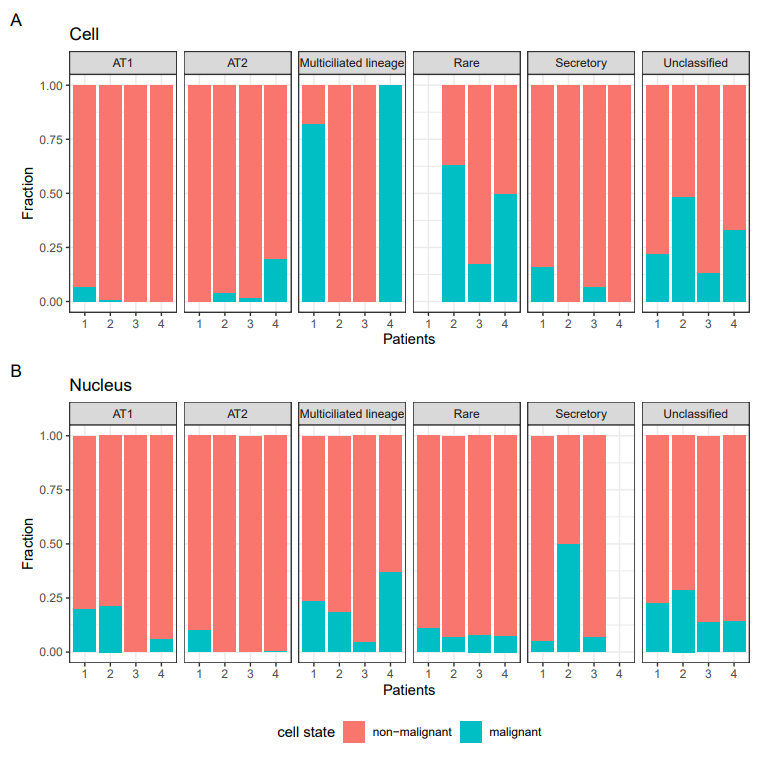

**S8 Fig |** The percentage of epithelial cells classified as malignant for each patient in *Cell* and *Nucleus* samples.

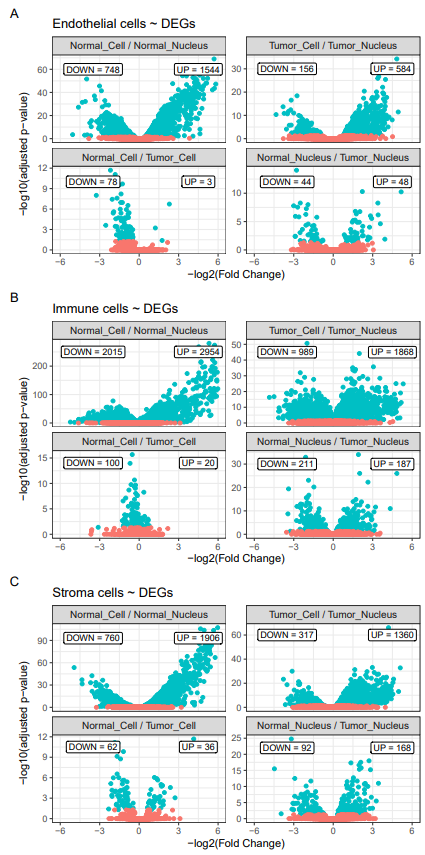

**S9 Fig |** DEGs (in turquoise) for Endothelial, Immune and Stroma cells with the number of up-regulated and down-regulated genes.

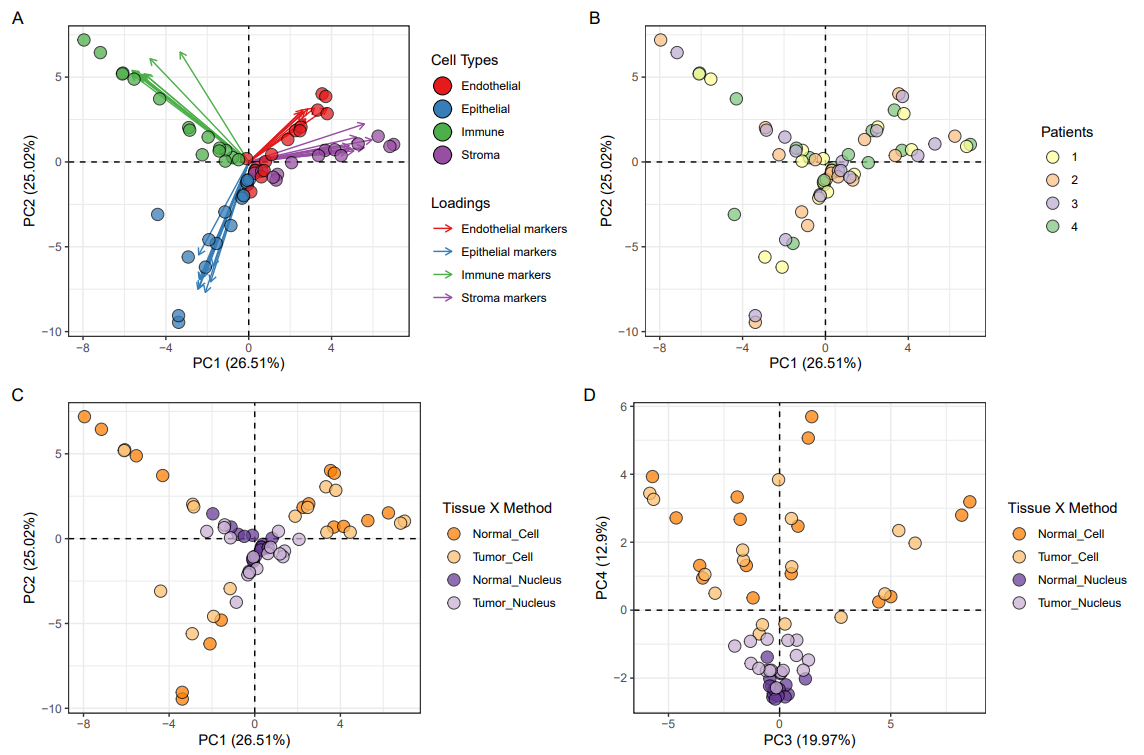

**S10 Fig | Principal Component Analysis** on the 39 marker genes used to distinguish between Immune, Epithelial, Endothelial and Stroma cell types (see Fig. S2 legend for a list of marker genes used). **A**. Marker genes loadings on the PCA (arrows colored by the cell type they are used to define) match well with the reference-based annotation of the samples (colored points). **B.** No bias in the clustering of the samples based on the patient identity. **C.** Samples cluster according to the method. Nucleus samples are closer to the center of the PCA, which implies that markers genes were less efficient in distinguishing between cell types in these samples. **D.** In Principal Components 3 and 4, Nucleus samples are separated by tissue type (Normal and Tumor).

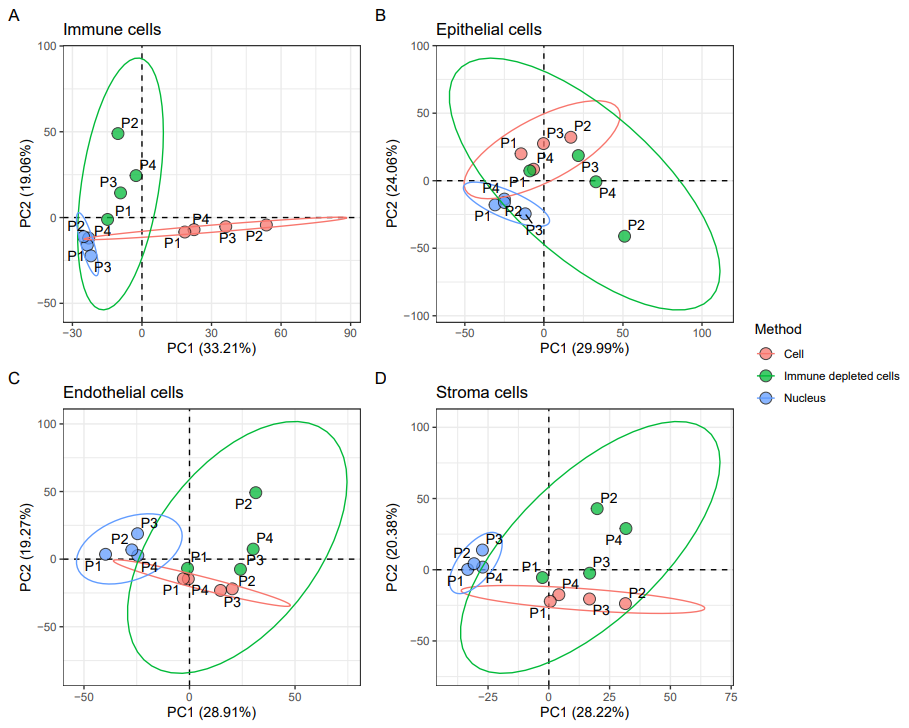

**S11 Fig | Principal Component Analysis** on the top 5 % most variable genes (Normal tissue) for **A.** Immune cells **B.** Epithelial cells **C.** Endothelial cells and **D.** Stroma cells. 95 % confidence interval ellipses are drawn for each method based on all four patients.
